# Microbiome composition modulates the lethal outcome of Drosophila A virus infection

**DOI:** 10.1101/2025.10.16.682821

**Authors:** Rubén González, Mauro Castelló-Sanjuán, Ottavia Romoli, Hervé Blanc, Hiroko Kobayashi, Jared Nigg, Maria-Carla Saleh

## Abstract

Host-associated microbiomes can strongly influence viral infection outcomes, yet how minor variations in commensal bacterial composition modulate viral pathogenesis remain poorly understood. Here, we used *Drosophila melanogaster* to investigate how bacterial microbiome composition affects pathogenesis of enteric RNA viruses. *Lactiplantibacillus plantarum* supplementation increased bacterial microbiome diversity without altering total bacterial load, while *Acetobacter pomorum* supplementation had minimal impact on the bacterial microbiome. *L. plantarum*-enriched flies exhibited an additional ∼15% reduction in lifespan from Drosophila A virus (DAV) infection despite showing reduced viral protein accumulation and similar viral RNA levels. The reduction in tolerance to viral infection required live bacteria and was observed only for DAV, as no change in mortality was observed with Nora virus or Drosophila C virus infections. Mechanistic investigations revealed that tolerance reduction occurs independently of transcriptional immune responses, as DAV-infected flies showed similar transcriptional profiles regardless of bacterial microbiome composition. Intestinal barrier function assays demonstrated that *L. plantarum*-supplemented flies died before developing signs of gut barrier disruption, suggesting that extra-intestinal mechanisms contribute to mortality; this interpretation is further supported by similar levels of intestinal damage markers observed in virus-infected flies under both microbiome conditions. Viral genomic sequencing ruled out microbiome-driven selection of more pathogenic viral variants, as no adaptive mutations were observed between microbiome conditions that could account for the differential pathogenesis. These findings describe how subtle shifts in microbiome composition modulate viral infection outcomes through pathways that operate independently of canonical immune responses, viral evolution, and intestinal damage.

## INTRODUCTION

Microbiomes shape host responses to viral infections across diverse biological systems (Fraune and Bosch, 2010; González and Elena, 2021; Karst, 2016; Kuss et al., 2011; Robinson and Pfeiffer, 2014; Pfeiffer and Virgin, 2016). However, dissecting the tripartite interactions between virus, host, and microbiome remains challenging due to microbiome complexity and limited experimental tractability.

*Drosophila melanogaster* provides an ideal system for such investigations. It is a simple, manipulatable model system that offers powerful mechanistic approaches to understand these tripartite interactions. The fly gut microbiome is remarkably simple, typically containing only 1-30 bacterial taxa (Broderick and Lemaitre, 2012; Storelli et al., 2011), in contrast to the complex diversity associated with vertebrates (>500 taxa) (Broderick and Lemaitre, 2012). However, the composition of fly bacterial microbiomes is strongly influenced by host diet and laboratory conditions. Indeed, gut-associated bacteria of Drosophila stocks can differ greatly between laboratories (Chandler et al., 2011, Broderick and Lemaitre, 2012). The most prevalent bacterial taxa in wild Drosophila are *Lactobacillales*, *Acetobacteraceae*, and *Enterobacteriaceae*, with most wild populations dominated by at least one of these (Adair et al., 2018; Chandler et al., 2011; McMullen et al., 2021). Additionally, *Lactobacillus*, *Acetobacter*, and *Enterococcus* are commonly reported in laboratory-reared flies (Broderick and Lemaitre, 2012; Ren et al., 2007; Cox and Gilmore, 2007). Within these dominant genera, natural populations commonly harbor multiple species: *Acetobacter* communities typically include *A. pomorum*, *A. tropicalis*, *A. orientalis*, and *A. persici*, while *Lactobacillus* communities are dominated by *L. plantarum* and *L. brevis*, with occasional presence of other species like *L. fructivorans* (Chandler et al., 2011; Staubach et al., 2013; Winans et al., 2017). *A. pomorum* is one of the most prevalent acetic acid bacteria in wild-caught flies and laboratory populations (Broderick and Lemaitre, 2012; Corby-Harris et al., 2007), and *Lactiplantibacillus plantarum* (formerly Lactobacillus plantarum, Zheng et al., 2020) is consistently identified as a key member of laboratory reared Drosophila microbiome, alongside other Lactobacillus species (Matos et al., 2017; McMullen et al., 2021; Wong et al., 2011). Both *L. plantarum* and *A. pomorum* have been found in most laboratory stocks analyzed across multiple studies (Broderick and Lemaitre, 2012) and have well defined impacts on host physiology; *Acetobacter pomorum* modulates metabolic homeostasis through insulin signaling (Shin et al., 2011), while *Lactiplantibacillus plantarum* promotes growth under nutrient limitation by enhancing protein assimilation (Storelli et al., 2011; Ryu et al., 2008). Together, the compositional and functional simplicity of the fly gut microbiome, combined with the ecological relevance of these dominant species, enables its precise and biologically relevant manipulation.

To elucidate the interaction between virus, host, and microbiome, we focused on Drosophila A virus (DAV), a single-stranded positive-sense RNA virus that is present in both natural and laboratory populations (Webster et al., 2015). DAV can be orally transmitted in laboratory settings, mimicking natural routes of infection (Nigg et al., 2021). DAV is an enteric virus that primarily infects the gut, which is the same niche occupied by commensal bacteria. Additionally, DAV infection disrupts intestinal homeostasis, reduces host lifespan, alters host locomotion, and accelerates host aging (Nigg et al., 2024; Castelló-Sanjuán et al., 2025; González et al., 2025). This infection phenotype presents a useful means to interpret how manipulating the host microbiome may impact the viral infection cycle and/or host responses to infection. Finally, DAV has a compact 5kb genome encoding only two proteins: an RNA-dependent RNA polymerase (RdRp) and capsid protein (CP), making it a tractable subject for evolutionary analysis.

In this study, we systematically dissected the three-way interactions between DAV, the *Drosophila* host, and its bacterial microbiome. We demonstrate that bacterial strain composition impacts viral pathogenesis (here defined as the detrimental impact of the infection in the host lifespan) through mechanisms that operate independently of transcriptional immune responses, intestinal barrier dysfunction, and viral adaptive evolution. Our findings reveal that host-microbiome-virus interactions can modulate disease tolerance through previously uncharacterized pathways with consequences for viral pathogenesis.

## RESULTS

### *L. plantarum* supplementation changes bacterial microbiome composition

We supplemented flies with bacteria to investigate how bacterial microbiome composition influences viral infection. We coated fly food with bacterial cultures starting one day before virus inoculation and continuing for five days post-inoculation, after which flies were maintained on standard food (Fig 1A, methods). To prevent direct bacteria-virus interactions outside the host, we performed viral inoculations in non-supplemented tubes before transferring flies to bacteria-supplemented food.

**Figure 1.**
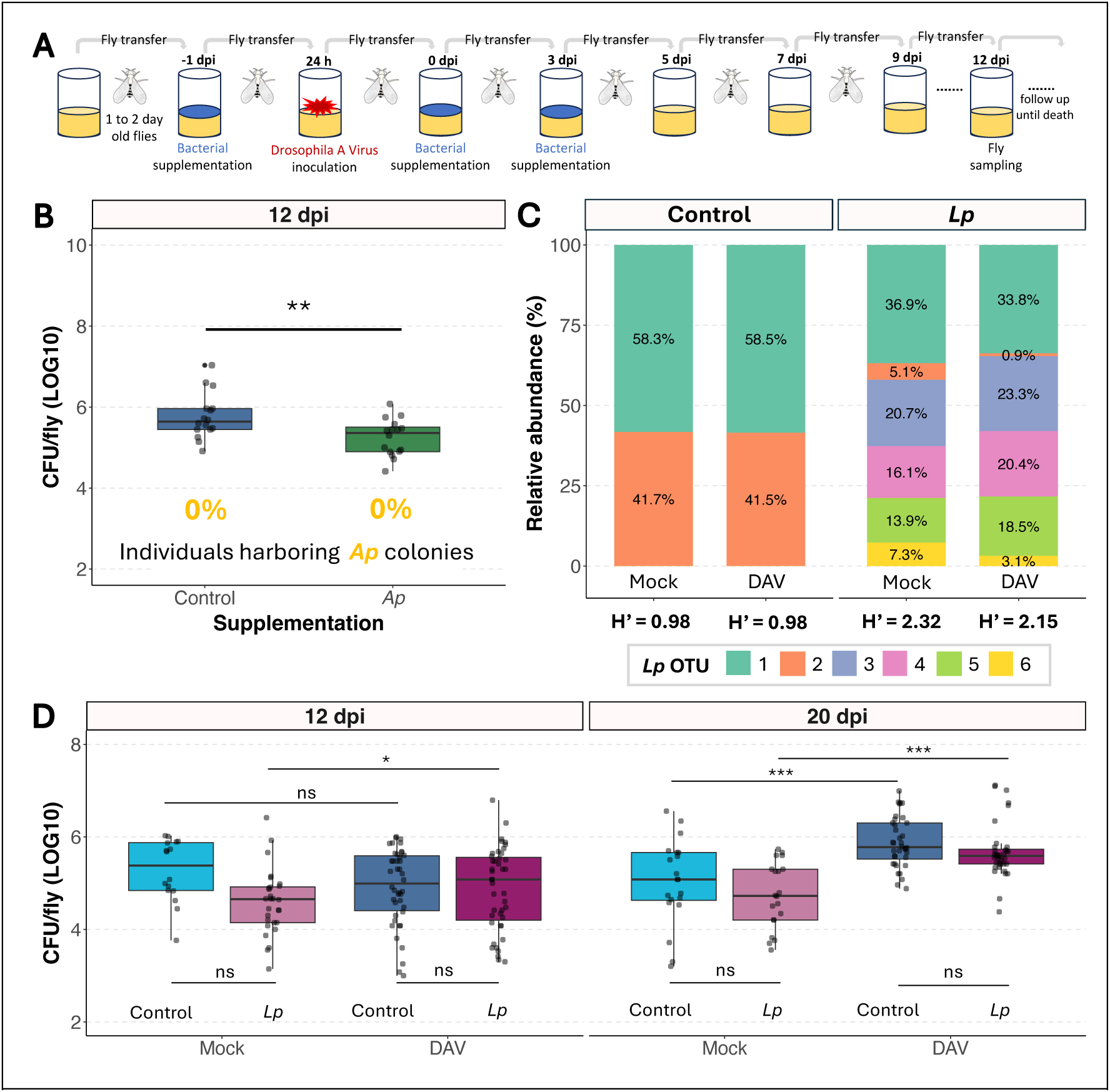
Bacterial microbiome characterization following bacterial supplementation. **A)** Experimental design schematic. **B)** Colony-forming units (CFU) per individual at 12 dpi in DAV-inoculated flies that were *Ap*-supplemented or not. Statistical comparison performed using a t-test on log₁₀-transformed data. **C)** Metagenomic analysis showing the microbiota composition of control versus *Lp-*supplemented flies. Stacked bars show relative abundance of the six different *L. plantarum* operational taxonomic units (OTUs), with each color representing a different OTU. Shannon diversity indices (H’) are displayed below each condition. Percentages within bars indicate relative abundance values**. D)** Total bacterial load (CFU/fly, log₁₀ scale) across treatment combinations at 12 and 20 dpi. Data show three experiments of Mock and DAV conditions for both control (non-supplemented) and Lp supplemented flies. Statistical analysis was performed using linear mixed-effects models with virus, treatment, and day as fixed factors and experiment as random factor, followed by pairwise comparisons using estimated marginal means with Tukey adjustment.

We first characterized how supplementation affected the fly gut bacterial microbiome. When bacterial microbiomes from *A. pomorum* WJL (*Ap*)-supplemented flies were plated 12 days post supplementation, we found no evidence of the characteristic yellow colonies that distinguish this bacterial species, and overall bacterial load was significantly lower compared to controls, indicating that in our conditions *Ap* fails to colonize the gut (Fig 1B). In contrast, *L. plantarum* WJL (*Lp*) supplementation substantially altered bacterial microbiome composition. *Lp* cultures produce a variable number of morphologically distinct rough-edged colonies when plated at 30° C (Supplementary Fig 1A), a colony morphology absent in the bacterial microbiome of our laboratory *w^1118^* fly stock. Following *Lp* supplementation, flies consistently harbored these characteristic rough-edged colonies that persisted well beyond the supplementation period and occurred independently of viral inoculation status (Supplementary Fig 1B). Long-read sequencing of bacterial 16S rDNA confirmed that only *L. plantarum* was detected in the flies, but different strains were detected in each condition: control flies (flies supplemented only with the broth media used to grow the bacterial cultures) harbored bacterial microbiomes dominated by two *L. plantarum* operational taxonomic units (OTUs), while *Lp*-supplemented flies contained more diverse *L. plantarum* OTU communities (Supplementary Fig 2). Shannon diversity (H’) analysis confirmed that *Lp* supplementation increased bacterial diversity (Fig 1C, Supplementary File 1). *Lp* supplementation increased bacterial strain diversity without altering total bacterial load (Fig 1D), though viral infection increased bacterial loads in all groups at later timepoints.

### Supplementation with live *Lp* reduces host tolerance to DAV infection

We orally inoculated flies with 1 OID₅₀ of DAV, the dose that infects 50% of exposed individuals. Infection rates were consistent (∼50%) across all treatments, indicating bacterial supplementation did not affect DAV infectivity (Fig 2A). However, *Lp* supplementation produced contrasting effects on viral accumulation and pathogenesis. *Lp*-supplemented flies showed reduced viral protein levels at 12 days post-infection (dpi) compared to control flies (Fig 2B), despite the viral RNA levels being similar at the same time point (Fig 2C). Analysis of viral RNA kinetics during early infection (0, 2, and 7 dpi) revealed similar virus accumulation patterns between control and *Lp*-supplemented flies in both gut and carcass tissues, with viral RNA appearing first in the gut and spreading systemically by day 2 (Supplementary Fig 3). This indicates that reduced protein levels were not due to altered viral dissemination or replication differences.

**Figure 2.**
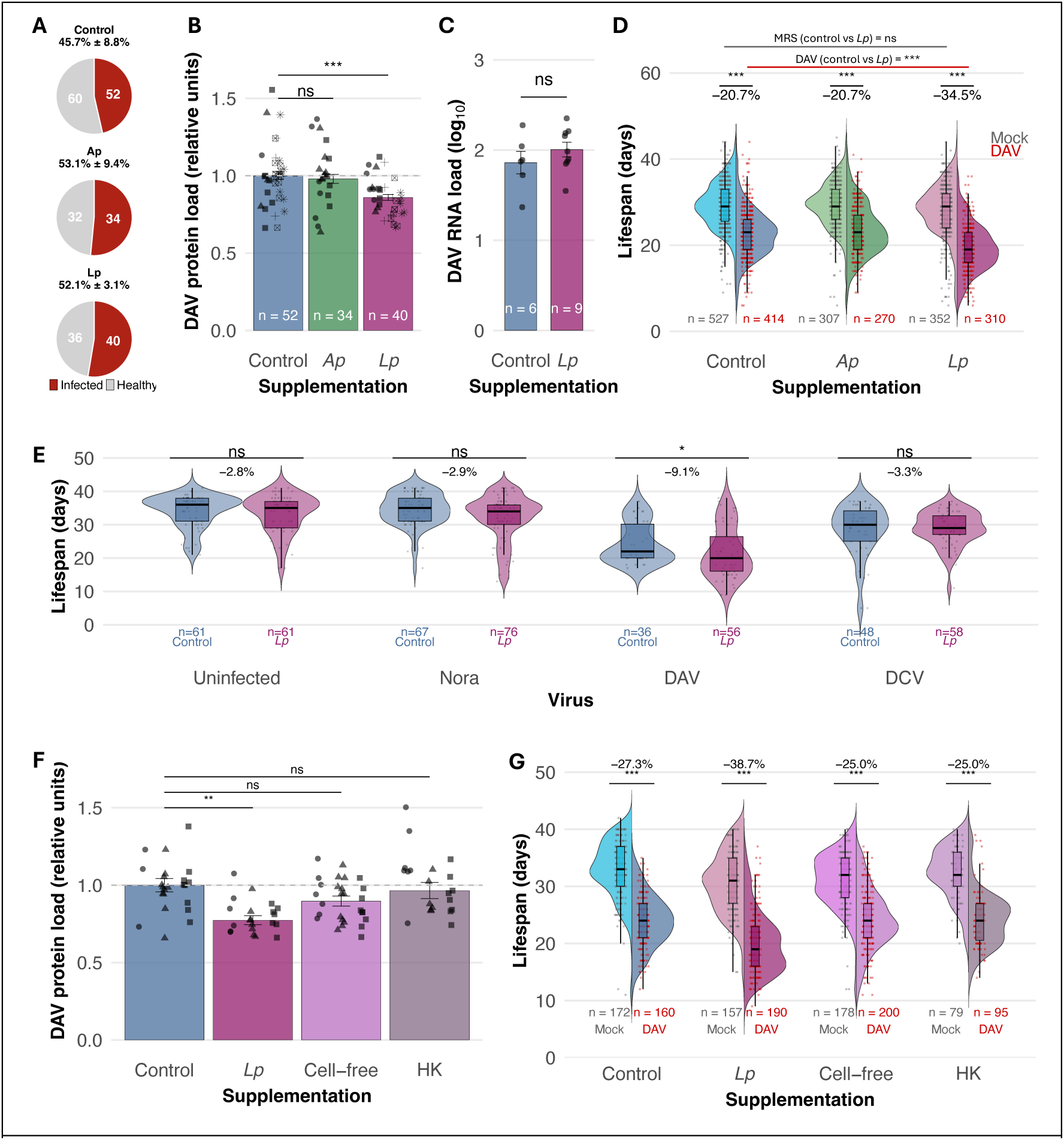
Differential impact of bacterial microbiome composition on viral infection dynamics. **A)** DAV infection prevalence in fly populations inoculated with 1 OID50 and supplemented with MRS (control), *A. pomorum* (*Ap*), or *L. plantarum* (*Lp*). **B)** Viral capsid protein accumulation across treatment groups. Significance was calculated using a general linear-mixed model where the supplementation was the fixed factor and the experimental block a random effect. Different shapes represent independent experiments. **C)** Comparative viral RNA accumulation in control versus *Lp*-supplemented flies. **D)** Lifespan distribution showing DAV-induced lifespan reduction across bacterial treatment groups. Significance was calculated using a general linear-mixed model where the supplementation and infection status were the fixed factors and the experimental block a random effect. Six experiments were performed with several tubes per condition tested in each experiment. **E)** Impact of *Lp*-enrichment on fly lifespan during infection with different persistent viruses. Significance was calculated using a general linear-mixed model where the supplementation was the fixed factor and the experimental block a random e:ect. **F)** Viral capsid protein accumulation in flies supplemented with control, viable *Lp*, cell-free *Lp* supernatant, or heat-killed *Lp*. Significance was calculated using a general linear-mixed model where the supplementation was the fixed factor and the experimental block a random effect. Different shapes represent independent experiments. One experiment was performed with several tubes per condition tested. **G)** Lifespan distribution of DAV-infected flies across *Lp* treatment conditions. Significance was calculated using a general linear-mixed model where the supplementation and infection status were the fixed factors and vial as a random effect. Two experiments were performed with several tubes per condition tested in each experiment.

Reduced viral protein levels did not, however, correlate with decreased pathogenesis. DAV infection in non-supplemented controls reduced fly lifespan by ∼20% compared to non-supplemented, mock-inoculated flies. DAV infection had a similar impact on lifespan in *Ap* supplemented flies compared to non-supplemented controls. In contrast, DAV-infected, *Lp*-supplemented flies showed ∼35% lifespan reduction compared to mock-inoculated, *Lp*-supplemented controls. This represents an additional ∼15% reduction compared to DAV-infected, non-supplemented flies (95% CI: 13.6% to 17.4%, *P* < 0.001). This difference between bacterial microbiome conditions was statistically significant when comparing DAV-infected flies, while no significant differences were observed between mock-infected flies across treatments (Fig 2D). Thus, *Lp* supplementation simultaneously reduced viral protein accumulation while enhancing virus-induced mortality.

To test whether the increased mortality observed after *Lp* supplementation was specific to a single-episode infections with DAV, we examined the effect of *Lp* supplementation in flies persistently infected with different RNA viruses. Similar to our observations with DAV single-episode infections, *Lp* supplementation drove increased mortality in flies harboring persistent DAV infections compared to persistently infected non-supplemented flies (Fig 2E). In contrast, flies persistently infected with Nora virus or Drosophila C virus (DCV) showed no survival changes under *Lp* supplementation, indicating that enhanced pathogenicity depends on DAV-specific interactions with the *Lp*-modified host environment.

We tested the bacterial requirements for modulating DAV pathogenesis using live *L. plantarum* cultures, cell-free supernatants, and heat-killed bacteria. Only viable bacteria reproduced both key phenotypes: reduced DAV protein accumulation (Fig 2F) and enhanced mortality (Fig 2G). Supernatants and heat-killed bacteria had no effect, demonstrating that viable bacteria are essential for modulating DAV pathogenesis.

### Reduction of DAV tolerance occurs independently of transcriptional immune responses

To investigate potential mechanisms underlying bacterial microbiome-mediated reduction of DAV tolerance, we examined host transcriptional responses. Prior to viral inoculation, flies with different bacterial microbiome compositions showed minimal transcriptional differences, with changes limited to genes related to nutrition and metabolism (Supplementary Fig 4).

We compared expression profiles of DAV-infected flies at 12 dpi to their respective mock-infected controls in both non-supplemented and *Lp*-supplemented flies. Both conditions mounted robust transcriptional responses to DAV infection, with hundreds of differentially expressed genes involved in immune defense, stress responses, and metabolic remodeling (Figure 3A, B). However, comparisons between the two bacterial microbiome conditions within each infection status revealed no significant differences between non-supplemented and *Lp*-supplemented flies in either mock-infected or DAV-infected states (Figure 3C, D). The absence of differentially expressed genes between bacterial microbiome conditions indicates that both non-supplemented and *Lp*-supplemented flies mount equivalent transcriptional responses to DAV infection.

**Figure 3.**
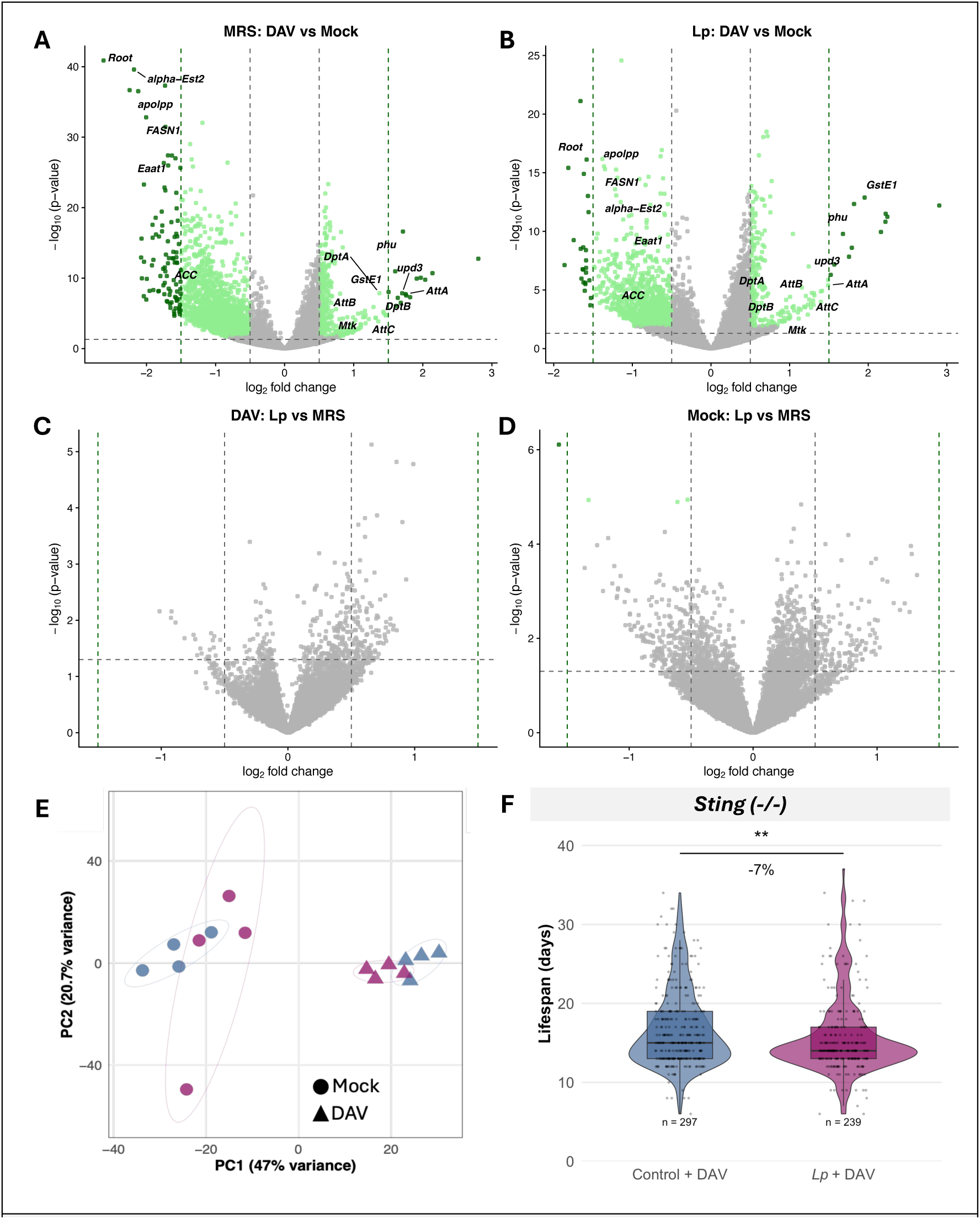
Transcriptional responses in each bacterial microbiome. **A-B)** Volcano plots showing differential gene expression in response to DAV infection (DAV-infected vs mock-infected flies) in non-supplemented (MRS) flies **(A)** and *Lp*-supplemented flies **(B)** at 12 dpi. The x-axis represents log₂ fold change and the y-axis represents -log₁₀ adjusted p-value. Horizontal dashed line indicates significance threshold (adjusted P < 0.05) and vertical dashed lines indicate fold change thresholds (|log₂FC| = 0.5). Green dots represent significantly differentially expressed genes, with darker green indicating higher magnitude changes (|log₂FC| > 1.5). Gray dots represent genes that do not meet significance criteria. Selected genes are labeled. Both bacterial microbiome conditions show similar transcriptional responses to DAV infection, with comparable numbers of differentially expressed genes and similar expression patterns. **C-D)** Direct comparison of gene expression between bacterial microbiome conditions. Volcano plots comparing *Lp-*supplemented versus non-supplemented flies in DAV-infected **(C)** and mock-infected **(D)** conditions. Horizontal dashed line indicates significance threshold (adjusted P < 0.05) and vertical dashed lines indicate fold change thresholds (|log₂FC| = 0.5). No genes show significant differential expression between bacterial microbiome conditions in either infected or uninfected flies, demonstrating that the reduced tolerance observed in *Lp-*supplemented flies occurs independently of transcriptional changes**. E)** Principal component analysis (PCA) of genome-wide gene expression profiles using variance-stabilized transformed counts. Each point represents a biological replicate (pool of 5 flies). Circles represent mock-infected samples and triangles represent DAV-infected samples. Blue indicates non-supplemented (MRS) conditions and purple/pink indicates *Lp*-supplemented conditions. **F)** Lifespan analysis of *Sting* mutant flies infected with DAV under different bacterial microbiome conditions. Violin plots show the distribution of individual fly lifespans, with horizontal lines indicating median values and quartiles. Each dot represents the lifespan of an individual fly. Statistical comparison performed using linear mixed-effects models with bacterial supplementation as fixed factor and vial as a random effect. Sample sizes are indicated on the plot. Each sample in panels A-D represents a pool of 5 flies (n = 4 biological replicates per condition).

Principal component analysis confirmed that infection status is the primary driver of transcriptional variance (PC1, 47%), while PC2 (20.7% variance) shows no clear separation by bacterial microbiome composition, confirming that the bacterial microbiome has a minimal influence on the fly’s transcriptional response (Figure 3E).

Given that transcriptional analysis cannot capture all aspects of immune function and considering the critical role of *Sting*-mediated innate immunity in DAV pathogenesis (Nigg et al., 2024), we tested whether this pathway mediates the bacterial microbiome-dependent reduction in viral tolerance. We supplemented *Sting* mutant flies with *Lp* and challenged them with DAV. *Lp*-supplemented *Sting* mutants exhibited approximately 10% additional reduction in lifespan compared to non-supplemented *Sting* mutants, similar to the effect observed in wild-type flies (Figure 3F). This demonstrates that the bacterial microbiome’s impact on host tolerance to DAV operates independently of Sting signaling, further supporting a mechanism that functions outside canonical antiviral immune pathways.

### Reduced DAV tolerance operates independently of intestinal pathology

To assess whether microbiome-mediated enhancement of DAV pathogenesis involves altered intestinal barrier function, we employed the "Smurf" assay, which uses a non-toxic blue dye to visualize gut barrier permeability (Rera et al., 2012). In mock-infected flies with control microbiomes, we observed the Smurf phenotype (indicating barrier disruption) in the days before death, establishing the normal progression of intestinal aging. As expected, DAV-infected flies with control microbiomes showed shorter lifespans and earlier appearance of Smurf phenotypes, with a reduced interval between barrier disruption and death (Figure 4A,B). This suggests that once the intestinal barrier is compromised, systemic effects accelerate mortality beyond what would be expected from intestinal damage alone.

**Figure 4.**
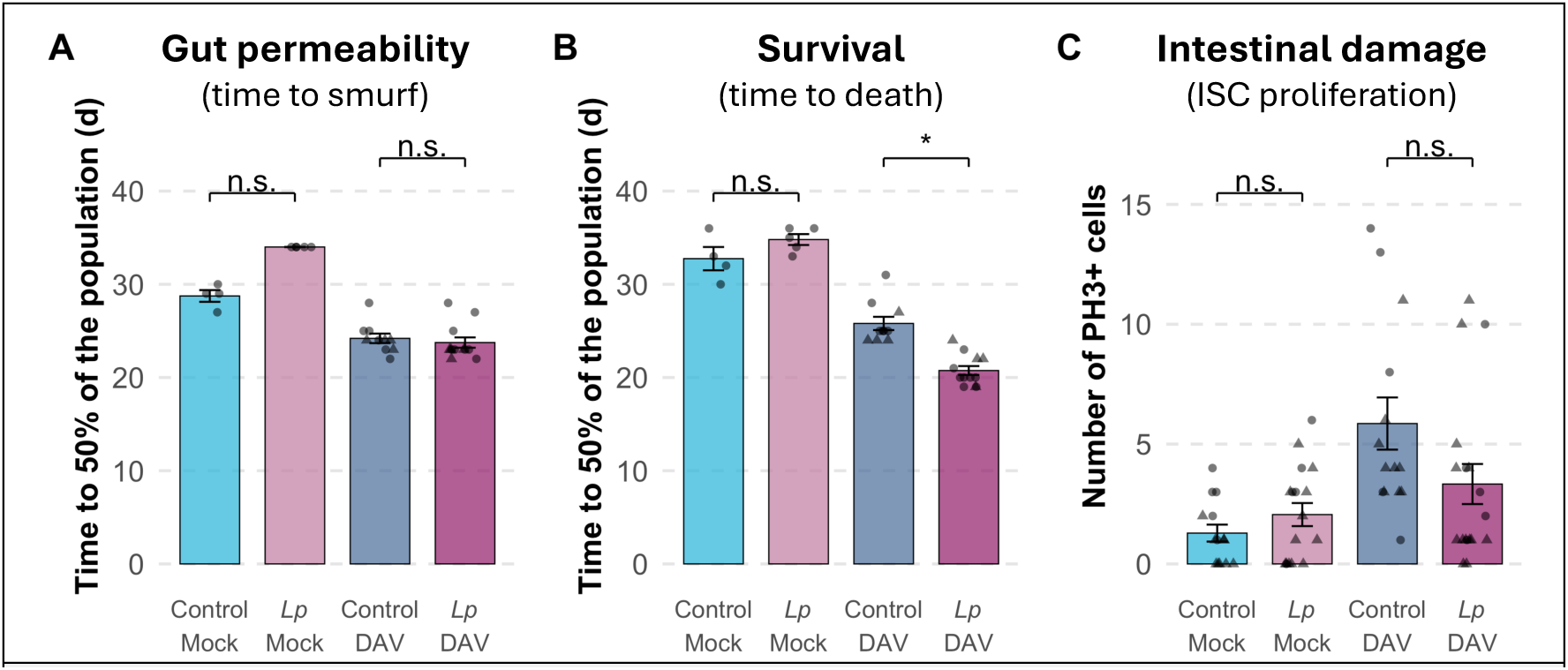
Reduced tolerance in *Lp*-supplemented flies operates through extra-intestinal mechanisms. **A-B)** Gut barrier permeability and survival measured in the same cohorts of flies. **(A)** Bars show the median time for 50% of the population to exhibit the Smurf phenotype (intestinal barrier disruption). DAV infection accelerates barrier disruption in both bacterial microbiome conditions. *Lp* supplementation does not significantly alter the timing of barrier disruption in either mock or DAV-infected flies**. (B)** Fly survival in the same experimental cohorts. DAV infection reduces lifespan in both conditions, with *Lp*-supplemented flies showing significantly greater lifespan reduction. **C)** Intestinal stem cell (ISC) proliferation quantified by PH3-positive cells in the midgut at 12 dpi. DAV infection significantly increases ISC proliferation compared to mock infection in both bacterial microbiome conditions, but Lp supplementation does not further enhance this proliferative response. This indicates that reduced tolerance in *Lp*-supplemented flies does not result from enhanced local intestinal damage. Data are presented as mean ± SE with individual data points shown. Statistical comparisons were performed using linear mixed-effects models with bacterial supplementation and infection status as fixed factors and vial as random effect. n.s., not significant; \**P* < 0.05

During DAV infection, *Lp*-supplemented flies exhibited the expected reduction in lifespan that we observed consistently across experiments. However, the median time to Smurf of *Lp*-supplemented and control-supplemented flies was similar during DAV infections (Figure 4A,B). This indicates that many flies died before showing signs of intestinal barrier disruption, suggesting that *Lp* supplementation reduces the host tolerance to DAV through mechanisms acting beyond the gut.

To further investigate intestinal effects, we quantified phospho-histone H3 (PH3)-positive cells in the midgut, a marker of intestinal stem cell (ISC) proliferation and epithelial damage response (Nigg et al., 2024). While DAV infection significantly increased the number of PH3-positive cells compared to mock infection, *Lp* supplementation did not further alter this proliferative response at 12 dpi (Figure 4C). This finding supports the observation that enhanced pathogenesis in *Lp*-supplemented flies does not result from increased local intestinal damage.

### Increased host mortality is not caused by the emergence of new viral variants

Given the observed differences in capsid protein accumulation and DAV pathogenesis between *Lp*-supplemented and non-supplemented bacterial microbiomes, we tested whether certain bacterial communities selected for more pathogenic viral variants during infection. To test this, we sequenced DAV genomes from flies with non-supplemented and *Lp*-supplemented bacterial microbiomes after 12 days of infection.

We identified 128 mutations across the viral genome, revealing three distinct evolutionary patterns: 45 mutations shared between conditions, 38 non-supplemented-specific, and 45 Lp-specific mutations (Supplementary Fig 5). Even within only 12 days, mutations with a notable frequency change (≥20%) were common in both conditions. Mutations were predominantly transitions over transversions and were distributed throughout the RdRp and capsid genes. Non-supplemented-specific mutations showed greater mean frequency increases (17.9 ± 3.8%) than *Lp*-specific ones (11.6 ± 1.1%), with a similar pattern in shared mutations (12.1 ± 3.2% increase in non-supplemented vs. 6.7 ± 1.9% in Lp). Over half of all mutations were novel, meaning they arose during the 12-day infection rather than being minor variants already present in the virus population used as inoculum. Despite this quasispecies diversity, no mutations reached fixation in any of the four biological replicates, and no adaptive mutations were identified that could account for the differential pathogenesis observed between microbiome conditions. These results demonstrate that while bacterial microbiome composition can influence viral population dynamics, the increased mortality in *Lp*-supplemented flies is not driven by selection of more pathogenic viral variants.

## DISCUSSION

Our study demonstrates that subtle bacterial microbiome alterations can reshape virus-host interactions through pathogen-specific mechanisms that operate independently of canonical immune transcription and adaptive viral evolution. *Lp* supplementation selectively enhanced DAV pathogenesis while not affecting survival during Nora virus or DCV infections, revealing the specificity of microbiome-virus interactions. This selectivity likely reflects diverse viral strategies in tropism, replication, and host response activation (Romano et al., 2022; Silva et al., 2021; Kuyateh and Obbard, 2023; Castelló-Sanjuán et al., 2025; Segrist et al., 2024). Similar virus-specific interactions occur in other systems: human norovirus requires specific bacteria for cell infection (Jones et al., 2014), while influenza pathogenesis varies with the bacterial species present (Ichinohe et al., 2011). The paradoxical reduction in viral protein levels alongside enhanced mortality indicates a decoupling of viral burden from pathogenesis, a pattern that distinguishes tolerance from resistance mechanisms. Resistance reduces pathogen load through immune-mediated elimination, while tolerance limits tissue damage and maintains physiological function despite infection (Simms and Triplett, 1994; Råberg et al., 2007). Our finding that *L. plantarum s*upplementation reduces host tolerance (ability to limit damage for a given pathogen load) rather than resistance (ability to limit pathogen burden) suggests that the microbiome modulates host responses to infection-induced damage rather than controlling viral replication directly. This distinction is critical because tolerance mechanisms can determine disease outcomes independently of pathogen burden and represent potential therapeutic targets that do not drive pathogen evolution (Lambrechts and Saleh, 2019). The observation that flies with lower viral protein levels nonetheless exhibited greater mortality exemplifies impaired tolerance: these flies were less capable of withstanding the physiological stress imposed by viral infection despite reduced viral burden.

Commensal bacteria influencing host-pathogen interactions is well-documented across diverse organisms (Hooper et al., 2012; Abt et al., 2012; Broderick et al., 2014; Belkaid and Harrison, 2017). However, our findings in Drosophila identify a tolerance-reduction mechanism that operates through non-canonical pathways. Three independent lines of evidence support this conclusion. First, transcriptional profiling revealed no gene expression differences between microbiome conditions, ruling out immune transcriptional responses. Second, intestinal pathology markers were similar across conditions, excluding gut damage as the driver. Third, the effect persisted in STING-deficient flies, showing independence a pathway critical for DAV pathogenesis.

Several non-mutually exclusive mechanisms warrant investigation. First, live *L. plantarum* may continuously produce metabolites that alter host cellular processes in ways that exacerbate viral pathology without changing steady-state gene expression. The requirement for viable bacteria (as heat-killed bacteria and supernatants had no effect) indicates that ongoing bacterial metabolism is essential. Second, bacterial enzymatic activities could modify host glycoproteins, lipids, or other biomolecules in ways that affect viral dissemination, cellular damage, or tissue-specific susceptibility. This possibility is exemplified by the *Serratia marcescens* protein that digests mucins in mosquito gut to enhance arboviral transmission (Wu et al., 2019). Third, post-transcriptional regulation could explain how microbiome composition affects pathogenesis without transcriptional signatures. Finally, the microbiome may alter host metabolic states that specifically affect cellular responses to DAV infection in ways that manifest only under the stress of viral replication.

Our viral genomic analysis revealed substantial mutational diversity arising within 12 days but no evidence that different microbiome conditions selected for more pathogenic viral variants. This supports that enhanced mortality results from altered host tolerance rather than viral adaptation to bacterial environments. The 12-day infection period may be sufficient to generate viral diversity but too short for adaptive mutations to arise, be selected, and reach frequencies that could drive the observed pathological differences. This finding contrasts with longer-term infections in other systems where microbiome composition can drive pathogen evolution (King et al., 2016; Ford et al., 2016).

Several limitations should be acknowledged when interpreting our results. First, while we have identified a reproducible and biologically significant phenotype, the underlying mechanism remains unresolved. Our data effectively rule out transcriptional immune responses and viral evolution as primary drivers, but the actual molecular and cellular mechanisms require further investigation. Second, we focused on bacterial components of the microbiome; fungal and archaeal community members may also contribute to the observed effects. Third, our study examined only female flies at a single temperature (29°C); sex-specific differences and temperature-dependent effects may exist. Fourth, we examined a limited number of viruses; the generalizability of these findings across different viral families remains to be determined.

Our results establish a novel phenotype in microbiome-virus interactions and provide a foundation for future mechanistic investigations. The finding that bacterial microbiome composition can significantly alter host tolerance to viral infection independently of immune transcriptional responses, viral evolution, and intestinal pathology opens new avenues for understanding how commensal bacteria influence disease outcomes.

## MATERIAL AND METHODS

### Animal strains and maintenance

We used mated female *w^1118^* flies for all experiments unless otherwise specified. The *Sting* mutant line contains the dSTING*^Rxn^* mutation and was a gift from J.L. Imler (Université de Strasbourg, France). We maintained all fly stocks on a cornmeal diet; each tube had approximately 7 mL of fly food, that was prepared by combining 440g inactive dry yeast, 440g corn meal, and 60g agar in 6L osmotic water. The mixture was autoclaved and after cooling, 150ml moldex solution (20% methylhydroxybenzoate) and 29ml propionic acid were added as preservatives. Flies were kept at 25°C under a 12:12 hour light:dark cycle and transfer to 29°C upon eclosion to perform the experiments. All stocks were verified to be free of *Wolbachia* infection.

### Bacterial culture

Flies were supplemented with bacteria by coating the fly food surface with 10^8^ CFU of a bacterial culture or MRS as a control (non-supplemented). We cultured *A. pomorum* strain WJL in 10 ml of MRS broth in 20 ml flasks at 30°C with 180 rpm agitation for 20 hours. We grew *L. plantarum* strain WJL in 10 ml of MRS broth (Carl Roth, Germany) in 15 ml culture tubes at 37°C without agitation for 20 hours. For CFU quantification, we plated serial dilutions on MRS agar and incubated plates for 48 hours at 30°C or 24 hours at 37°C.

To prepare cell-free supernatant, we pelleted 2 ml of a 10^9^ CFU/ml bacterial culture by centrifugation at 5000 rpm for 10 minutes and filtered the supernatant through a 0.22 μm filter. For heat-killed cultures, we incubated 1 ml of 10^9^ CFU/ml culture at 100°C for 10 minutes. We verified the absence of viable cells in both preparations by plating 50 μl on MRS agar and confirming no colony growth after 48 hours of incubation at 30°C.

### Viruses

We orally inoculated mated female flies 1-2 days post-eclosion. After maintaining males and females together at 29°C for 24 hours, we removed males and transferred females to empty vials for three hours of starvation. We then moved flies to vials containing food coated with 100 μl of DAV stock (1-2 OID₅₀) and maintained groups of 20-30 flies per vial for 24 hours. We designated the transfer time to fresh vials as 0 dpi and transferred flies to new vials every 2-3 days thereafter. Mock inoculations followed the same procedure using filtered extract from uninfected flies.

For persistent infections, we used persistently infected flies from Nigg and colleagues (2024). The stocks were maintained at 25°C under a 12:12 light:dark cycle.

### Preparation of viral inoculum

We prepared DAV stocks from persistently infected w1118 flies following Nigg et al. (2024). We homogenized 60 mixed-age flies in 300 μl PBS, snap-froze the homogenate, and clarified it by two rounds of centrifugation (15,000 × g, 10 minutes, 4°C). After filtering through a 0.22 μm filter, we prepared 100 μl aliquots for storage at -80°C. We prepared mock stock identically using uninfected flies.

We determined viral titer via 50% endpoint dilution (Nigg et al., 2021), using ELISA to assess infection status at 12 dpi in flies exposed to serial dilutions of viral stock. For experiments, we used 1:5 dilutions from the stock for the 1000 OID inoculum and 1:500 for the 1 OID inoculum.

### Enzyme-linked immunosorbent assay (ELISA)

We detected DAV infection by ELISA following Nigg et al. (2021). We homogenized individual flies in 100 μl PBS and mixed 20 μl homogenate with 20 μl lysis buffer (40 mM HEPES pH 7.5, 2 mM DTT, 200 μM KCl, 10% glycerol, 0.1% NP-40, 1× complete EDTA-free Protease Inhibitor Cocktail (Roche, 11873580001). After 15 minutes at room temperature, we added 10 μl of this mixture to 190 μl of 0.05 M carbonate-bicarbonate buffer (pH 9.6) in MaxiSorp plates (Thermo Fisher, 442404) and incubated for 2 hours at room temperature.

We washed plates three times with PBS-T (1× PBS, 0.05% Tween-20), blocked with PBS-T containing 5% non-fat dry milk (2 hours at room temperature), and incubated overnight at 4°C with a polyclonal anti-DAV CP antibody (1:2000). After washing, we added donkey anti-rabbit IgG-HRP (Cytiva, NA934; 1:2000) for 2 hours at room temperature, washed, and developed with TMB substrate (Thermo Fisher, 34022). We stopped the reaction with 2N HCl and measured absorbance at 450 nm in a plate reader (Tecan Infinite M200 PRO). We defined infection as A₄₅₀ values exceeding the mock average plus three standard deviations.

For viral protein quantification, we used the absorbance values at 450 nm as a measure of viral capsid protein abundance. We normalized all values within each plate by dividing by the mean absorbance of DAV-positive non-supplemented samples on that plate. The normalized values are presented as viral capsid protein accumulation.

### Lifespan analysis

We conducted lifespan experiments using three to six biological replicates with 20-25 female flies per replicate, sorted at five days post-infection. We maintained flies at 29°C under the specific vial conditions described for each experiment. We monitored survival daily by counting dead flies in each vial and transferred flies to fresh vials every three to four days.

### Smurf assay

We assessed intestinal barrier function using the Smurf assay with biological replicates of 20 female w1118 flies, set up as described for survival analysis. Beginning at 7 dpi, we continuously maintained flies in vials containing food supplemented with 100 μl of sterile 16% FD&C blue dye. We scored "Smurfness" (blue dye leakage throughout the body) daily by examining flies according to Martins et al. (2018).

### RNA extraction

We extracted total RNA from whole flies by resuspending 100 μl of fly homogenate (in PBS buffer) in 400 μl TRIzol Reagent (Invitrogen, 15596026), vortexed thoroughly, and incubated for 3 minutes at room temperature. We added 100 μl chloroform, vortexed again, and incubated for another 3 minutes before centrifuging at 12,000 × g for 15 minutes at 4°C. We collected the upper aqueous phase and added an equal volume of 100% isopropanol and 1 μl of GlycoBlue (ThermoFisher Scientific), vortexed thoroughly, and incubated for 3 minutes at room temperature. We then centrifuged at 12,000 × g for 10 minutes at 4°C. After discarding the supernatant, we washed the RNA pellet with 500 μl of 75% ethanol, inverted 10 times to mix, and centrifuged at 7,500 × g for 5 minutes at 4°C. We removed the supernatant, air-dried the tube, dissolved the RNA pellet in 20 μl Milli-Q water, and incubated at 55°C for 10 minutes. We measured RNA concentration using the Qubit RNA BR Assay Kit (Invitrogen, Q10211).

### Immunofluorescence of digestive tissues

We dissected whole digestive tracts in 1× PBS over a 20-minute period and immediately fixed them in 4% paraformaldehyde for an additional 50 minutes. We washed fixed tissues three times (10 minutes each) in 1× PBS with 0.1% Triton X-100 (PBT), incubated them in 1× PBS with 50% glycerol for 30 minutes, and equilibrated them in 1× PBT for 30 minutes. We then incubated tissues with anti-PH3 primary antibody (rabbit, 1:1000, Merck Millipore, 06-570) diluted in 1× PBT overnight at 4°C. After three 10-minute washes, we incubated tissues with anti-rabbit Alexa Fluor 647 secondary antibody for 3-5 hours at room temperature. We performed three final 10-minute washes, with the last wash containing 1 μg/ml DAPI, and mounted tissues in 4% N-propyl-gallate in 80% glycerol. We imaged samples using a Zeiss LSM 700 confocal microscope at the Institut Pasteur Photonic BioImaging facility and manually counted PH3-positive cells throughout the entire midgut.

### Quantitative reverse transcription polymerase chain reaction (RT-qPCR)

We performed reverse transcription on total RNA using random primers and Maxima H Minus Reverse Transcriptase (Thermo Scientific, EP0751) according to the manufacturer’s instructions. We diluted the resulting cDNA 1:10 with nuclease-free water and performed qPCR in triplicate 10 μl reactions using virus-specific primers (Supplementary Table 1). The thermal cycling protocol consisted of 2 minutes at 50°C, 10 minutes at 95°C, followed by 40 cycles of 15 seconds at 95°C and 60 seconds at 60°C, with a standard melt curve analysis after amplification. We set the threshold for viral RNA detection at Ct 35, considering samples with higher Ct values as uninfected. We normalized virus Ct values against the Rp49 housekeeping gene and presented results as log10 of 2^-ΔCt^ values.

### High-throughput sequencing

We performed RNA sequencing on 4 biological replicates per experimental condition. We collected flies at 12 dpi from four treatment groups (control+mock, control+DAV, Lp+mock, Lp+DAV) and also sequenced samples from both supplementation conditions immediately before virus inoculation. We confirmed infection status by ELISA and prepared 4 pools of 5 individuals per condition. We prepared RNA-seq libraries from 15 ng of pooled RNA (3 ng per individual) using the NEBNext Ultra II RNA Library Prep Kit (New England Biolabs, E7770L) with NEBNext Multiplex Oligos (Dual Index Primers Set 1; New England Biolabs, E7600S). We performed all sequencing on an Illumina NextSeq 500 instrument using a NextSeq 500/550 High Output Kit v2.5 (75 cycles; Illumina, 20024906).

### Bacterial microbiome composition sequencing

We collected 60 female flies per condition at 12 dpi. We surface-sterilized flies to remove external microbes through sequential washes: 10% bleach for 5 minutes, 70% ethanol for 5 minutes, followed by three rinses in sterile water. We added lysis buffer (10 mM Tris-HCl pH 8, 26 mM EDTA, 0.5% SDS, 5 mg/ml lysozyme) to the flies and then homogenized with sterile pestles. After incubating lysates for 1 hour at 37°C, we extracted total DNA using the DNeasy Blood and Tissue Kit (Qiagen) following the manufacturer’s protocol.

We amplified the 16S rRNA gene using 100 ng genomic DNA as template in 50 μl final reaction volume. We employed 27F and 1391R primers (Supplementary Table 1) and performed PCR using Phusion High Fidelity Polymerase (Thermo Fisher), using 1 U of enzyme and a final primer concentration of 400 nM. The PCR cycle was: an initial denaturation step of 5 min at 98 °C followed by 35 cycles of 98 °C for 15 sec, 60 °C for 20 sec and 72 °C for 1 min, followed by a final extension at 72 °C for 10 min. PCR amplicons were purified using the NucleoSpin gel and PCR clean-up purification kit (Macherey Nagel) and sequenced by Plasmidsaurus using Nanopore. Between 28522 and 51439 reads were obtained per sample.

We quality-checked the resulting long-reads using NanoPlot (version 1.42.0) and filtered them with filtlong (version 0.2.1), discarding reads with average quality scores below Q17 and read lengths outside the 1200-1500 bp range. Reads were then denoised and chimeras were removed using dada2 in qiime2 (version 2019.10). OTU taxonomy was assigned using the Silva database.

### Genome analysis

We trimmed 15 nucleotides from the 5ʹ end of raw sequencing reads and removed reads with quality scores below 30. We mapped the clean reads to the *Drosophila melanogaster* genome (release dmel-r6.56) using STAR version 2.7.11b (Dobin et al., 2013). We performed feature counting with HTSeq version 0.11.2 (Putri et al., 2022) and differential gene expression analysis with DESeq2 in R Studio (version 2023.03.1). We filtered genes with fewer than 10 total counts across all samples and applied log2 fold change shrinkage using the "normal" method. We identified differentially expressed genes between DAV-infected and mock-inoculated flies within each bacterial microbiome condition and performed variance stabilizing transformation for visualization. We considered only adjusted p-values for statistical significance.

For viral genomic analysis, we first sequenced our DAV inoculum and mapped it to a reference DAV genome (GenBank accession no. FJ150422.1) to establish a stock-specific reference sequence. We processed experimental samples by filtering reads shorter than 40 nucleotides using cutadapt and trimmed 17 nucleotides from the 5’ end of reads. We aligned the processed reads to the reference genome using HiSAT2 version 2.2.1 with parameters optimized for viral sequences (--no-spliced-alignment, --score-min L,0.0,-0.8). We converted alignments to sorted BAM format using samtools and assessed viral load using samtools *flagstat*. For variant detection, we applied indel quality correction using LoFreq *indelqual*, followed by variant calling with LoFreq using a minimum coverage threshold of 5 reads and significance level of 0.05.

### Statistical analysis

We performed all statistical analyses in R version 4.3.2 using the Rstudio development environment version 2023.12.1+402. Statistical significance was defined as adjusted *P* < 0.05, with symbols representing significance levels as follows: *** *P* < 0.001; ** *P* < 0.01; * 0.01 < *P* < 0.05.

We applied generalized linear models (GLM) using the glm function from the "stats" package and linear mixed-effects models (LMM) using the lmer function from the "lme4" package version 1.1-35.5, selecting the appropriate model based on data structure and experimental design. For post-hoc analyses, we conducted pairwise comparisons using the *emmeans* function from the "emmeans" package version 1.10.5, applying Tukey’s correction to control for multiple testing. Detailed descriptions of the specific statistical approaches used for each analysis are provided in the corresponding figure legends.

## ACKNOWLEDGEMENTS

We thank François Leulier laboratory for sharing the bacterial strains. We thank members of the Saleh lab for fruitful discussion, Cassandra Koh for assistance with experiments, and Anamarija Butković for guidance in the phylogenetic analysis.

## AUTHOR CONTRIBUTIONS

**Rubén González:** conceptualization, data curation, formal analysis, investigation, visualization, writing-original draft, writing-review, supervision, and editing. **Mauro Castelló-Sanjuán:** investigation. **Ottavia Romoli:** investigation, formal analysis, writing-review, and editing. **Hervé Blanc:** investigation. **Hiroko Kobayashi:** investigation. **Jared Nigg:** writing-review, supervision. **Maria-Carla Saleh:** conceptualization, funding acquisition, project administration, resources, supervision, writing-review, and editing. All authors gave final approval for publication.

## DATA AVAILABILITY

Raw sequencing reads can be found in the Sequence Read Archive under the BioProject PRJNA1293387. All other data supporting the findings are available within the article and supplementary files.

## SUPPLEMENTARY MATERIAL

**Supplementary Figure 1.**
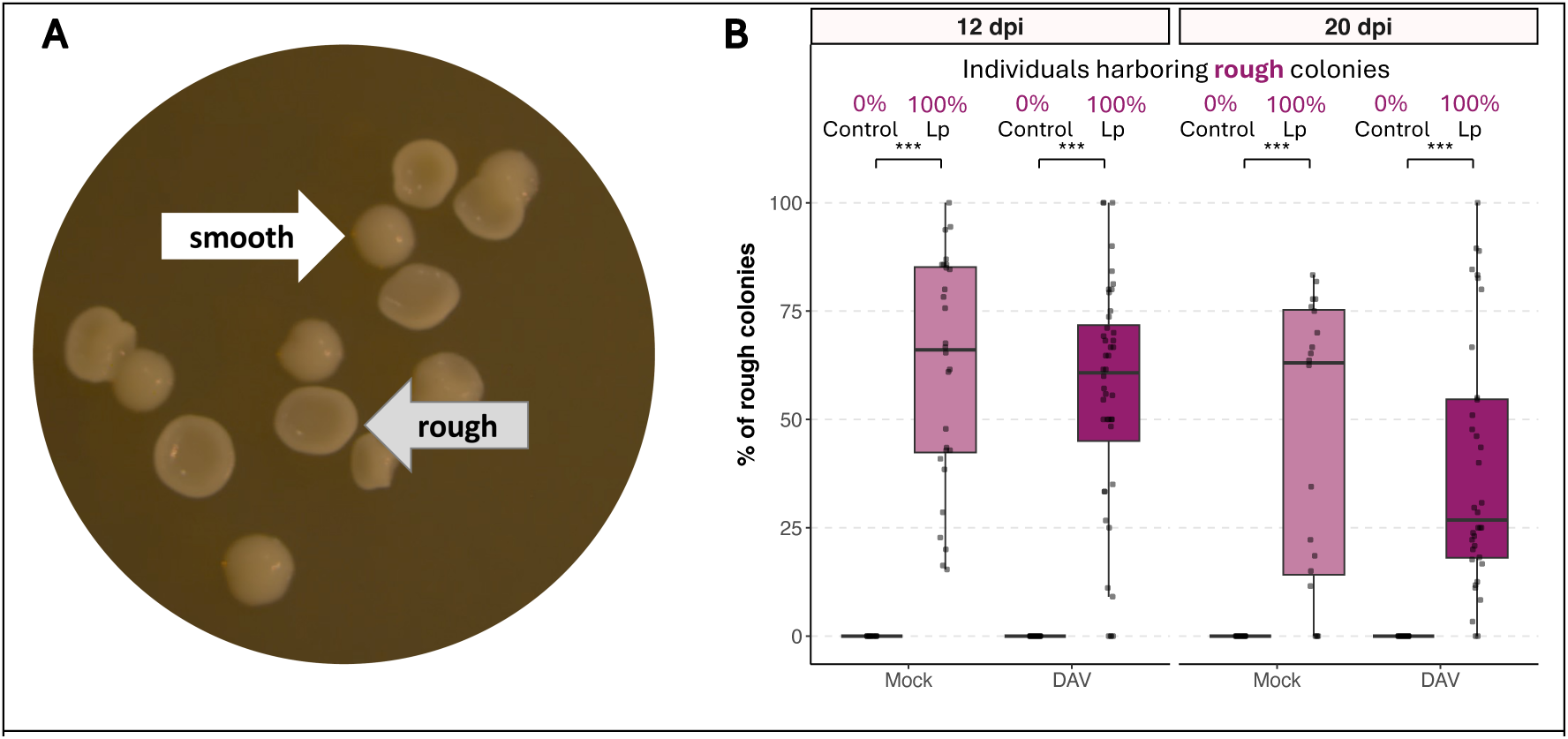
**A)** Morphologically distinct rough colonies in *L. plantarum* WJL cultures that appear when plated at 30 °C. **B)** Quantification of rough colony prevalence and abundance in control versus Lp-supplemented flies at 12 and 20 dpi.

**Supplementary Figure 2.**
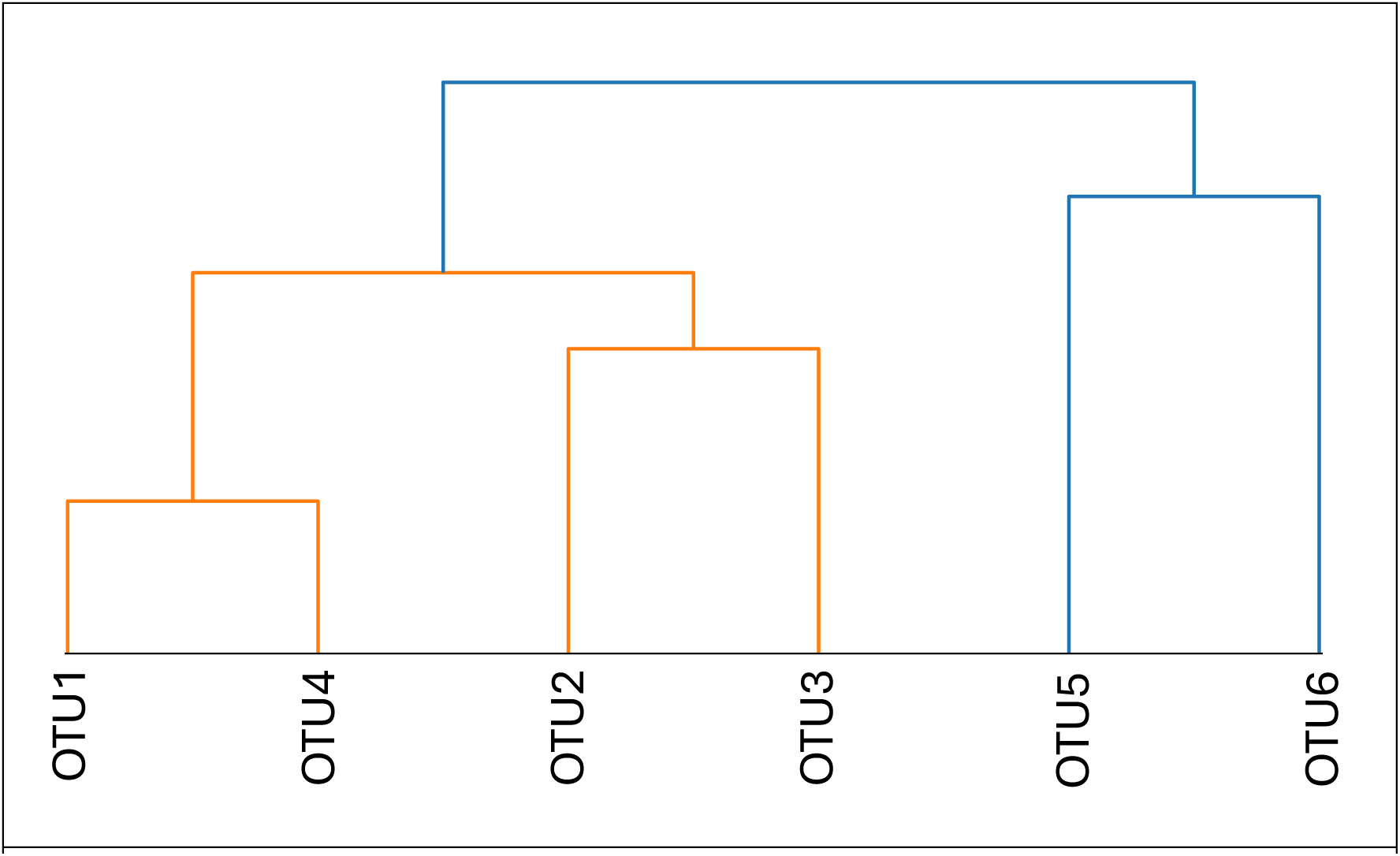
Hierarchical clustering of *L. plantarum* OTU 16S rDNA sequences. Multiple sequence alignment of bacterial 16S rDNA sequences was performed using MAFFT v7. Pairwise genetic distances were calculated between aligned sequences, and hierarchical clustering using average linkage was performed to visualize sequence similarity relationships among the six *L. plantarum* OTUs detected in fly bacterial microbiomes. Branch lengths represent genetic distance (proportion of differing nucleotide positions).

**Supplementary Figure 3.**
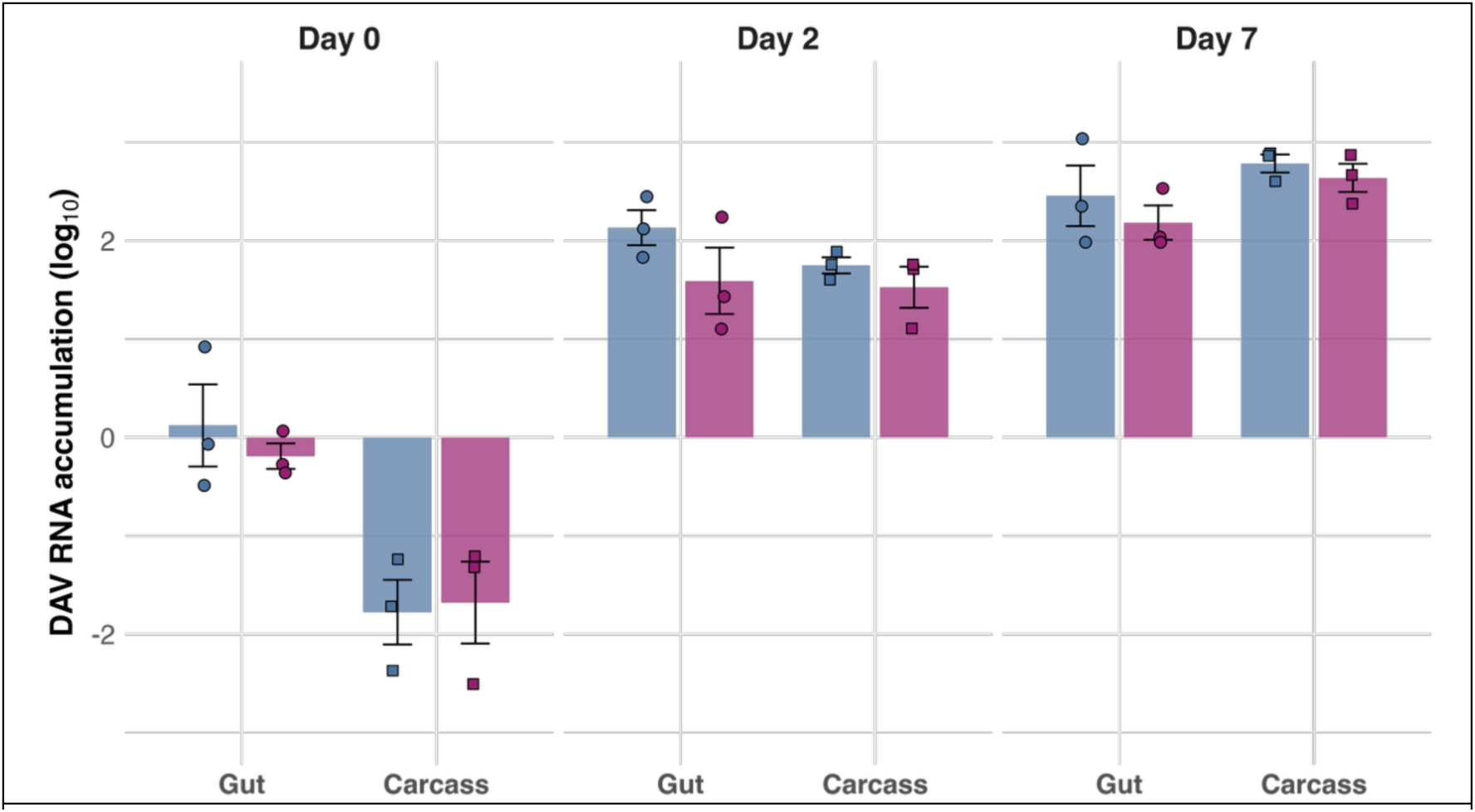
DAV RNA accumulation in gut and carcass tissues following oral inoculation. DAV RNA levels were measured in gut and carcass samples at 0, 2, and 7 days post-inoculation with 1000 OID of DAV and expressed as log₁₀ accumulation values. Blue bars represent non-supplemented flies and purple bars represent Lp supplemented flies. Individual data points are shown as circles (gut samples) and squares (carcass samples), with each point representing a pool of 5 individuals (n = 3 pools per group per time point). Data are presented as mean ± SE.

**Supplementary Figure 4.**
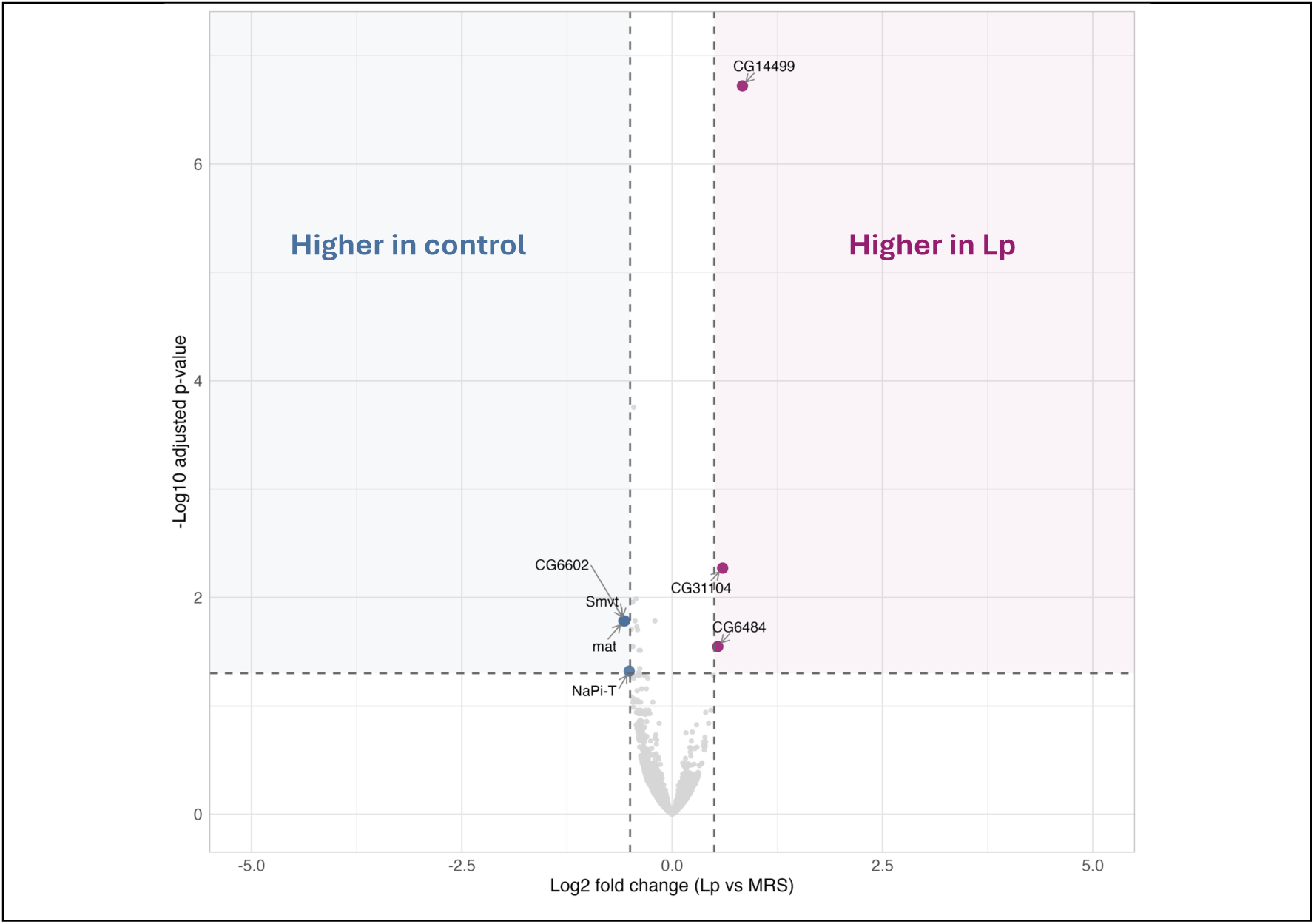
Transcriptional differences between bacterial microbiome conditions prior to viral inoculation. Volcano plot showing differential gene expression between *Lp-*supplemented and control (non-supplemented) flies before DAV inoculation. The x-axis represents log₂ fold change (*Lp* vs control) and the y-axis represents -log₁₀ adjusted p-value. Horizontal dashed line indicates significance threshold (adjusted *P* < 0.05) and vertical dashed lines indicate fold change thresholds (±0.5 log₂ fold change). Gray dots represent genes that do not meet significance criteria. Colored dots represent significantly differentially expressed genes, with blue indicating genes more highly expressed in control flies and purple indicating genes more highly expressed in Lp-supplemented flies. Significant genes are labeled with their names.

**Supplementary Figure 5.**
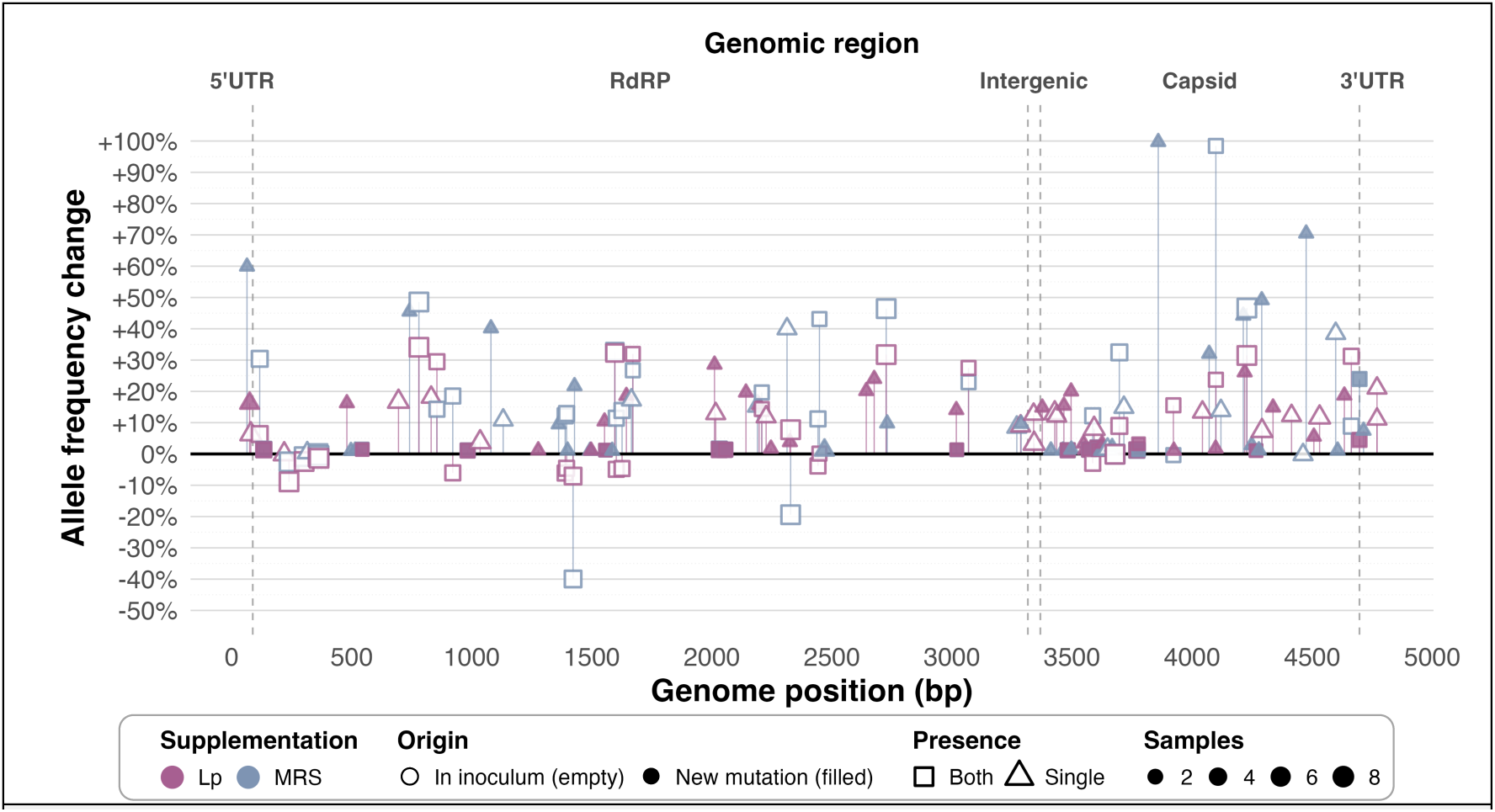
Viral genomic diversity across microbiome conditions. Allele frequency changes across the DAV genome comparing viral populations from control and *Lp-*supplemented flies at 12 days post-infection. The x-axis represents genomic position (bp) with major genomic regions indicated above, and the y-axis shows percent change of mutation frequency relative to the viral stock used as inoculum. Blue symbols indicate mutations in non-supplemented conditions while purple symbols indicate mutations in *Lp-*supplemented conditions. Squares represent mutations present in both conditions and triangles represent bacterial microbiome-specific mutations. Empty symbols indicate mutations already present in the inoculum and filled symbols indicate newly emerging mutations. Symbol size reflects the number of samples in which each mutation was detected. Each sample represents a pool of 5 individual flies and each condition has 4 samples.

**Supplementary Table 1.**
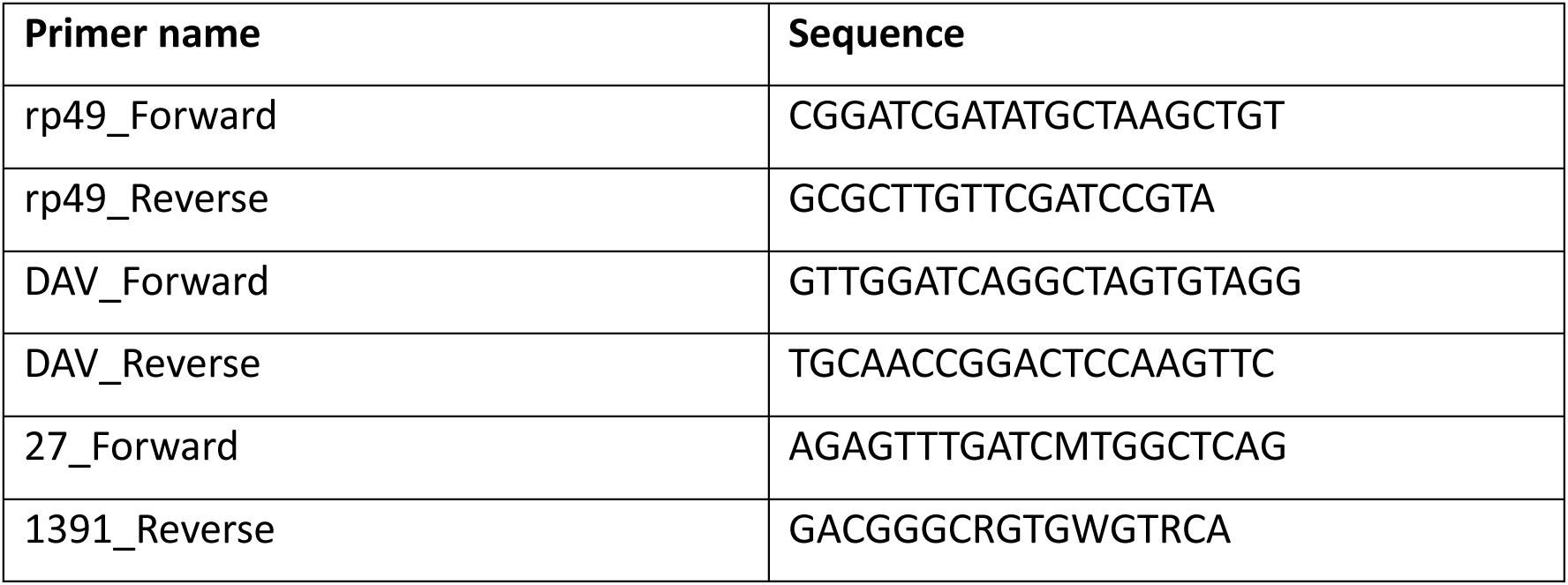
Primers used in qPCR step of RT-qPCR.

**Supplementary File 1. Bacterial operational taxonomic unit (OTU) sequences and taxonomic assignments.** FASTA file containing consensus sequences for all six *Lactobacillus plantarum* OTUs identified through 16S rDNA sequencing.

**Supplementary File 2. RNA-seq mapped read counts for all samples**. Excel file containing raw mapped read counts for all genes across all experimental conditions. Data include four biological replicates (pools of 5 flies each) for each of the following conditions: non-supplemented mock-infected, non-supplemented DAV-infected, *L. plantarum*-supplemented mock-infected, and *L. plantarum-*supplemented DAV-infected flies at 12 days post-infection, as well as samples from both supplementation conditions immediately before viral inoculation. Read counts were generated using HTSeq version 0.11.2 following alignment to the *Drosophila melanogaster* genome (release dmel-r6.56) with STAR version 2.7.11b as described in Methods.

**Supplementary File 3. DAV genome mutations and frequency changes from the viral stock used for inoculation.** Excel file containing comprehensive analysis of viral mutations detected in DAV populations from non-supplemented and *Lp*-supplemented flies at 12 dpi.

**Supplementary File 4. Source data for all figures.** Excel file containing individual data points and measurements for all main and supplementary figures. Each worksheet corresponds to one figure panel with raw values, sample sizes, and replicate information.

